# Impact of pre-anthesis drought stress on physiology, yield-related traits and drought responsive genes in green super rice

**DOI:** 10.1101/2021.11.18.469071

**Authors:** Hassaan Ahmad, Syed Adeel Zafar, Muhammad Kashif Naeem, Sajid Shokat, Safeena Inam, Amir Shahzad Naveed, Jianlong Xu, Zhikhang Li, Ghulam Muhammad Ali, Muhammad Ramzan Khan

## Abstract

Optimum soil water availability is vital for maximum yield production in rice which is challenged by increasing spells of drought. The reproductive stage drought is among the main limiting factors leading to the drastic reduction in grain yield. Objective of this study was to investigate the molecular and morpho-physiological responses of pre-anthesis stage drought stress in green super rice. The study assessed the performance of 26 rice lines under irrigated and drought conditions. Irrigated treatment was allowed to grow normally while drought stress was imposed for 30 days at pre-anthesis stage. Three important physiological traits including pollen fertility percentage (PFP), cell membrane stability (CMS) and normalized difference vegetative index (NDVI) were recorded at anthesis stage during the last week of drought stress. Agronomic traits of economic importance including grain yield were recorded at maturity stage. The analysis of variance demonstrated significant variation among the genotypes for most of the studied traits. Correlation and principal component analyses demonstrated highly significant associations of particular agronomic traits with grain yield, and genetic diversity among genotypes, respectively. Our study demonstrated a higher drought tolerance potential of GSR lines compared to local cultivars, mainly by higher pollen viability, plant biomass, CMS, and harvest index under drought. In addition, the molecular basis of drought tolerance in GSR lines was related to upregulation of certain drought responsive genes including *OsSADRI, OsDSM1*, and *OsDT11*. Our study identified novel drought tolerance related genes (*OsDRG-1, OsDRG-2, OsDRG-3* and *OsDRG-4)* that could be further characterized using reverse genetics to be utilized in molecular breeding for drought tolerance.

## INTRODUCTION

Rice (*Oryza sativa* L.) is one of the primary staple food crops for nearly 50% of the world population (Zafar *et al*. 2018). The countries located in East Asia, South Asia, and South East Asia are dominant in production and consumption of rice across the globe. Historically, more than 90 percent of world rice production is contributed from these countries (Barker *et al*. 1999). It’s production is needed to increase by 0.6 to 0.9% per year till 2050 to feed the further 2 billion people (Desa 2015). However, different abiotic and biotic stresses are major limiting factors for obtaining higher yield in rice (Zafar *et al*. 2017; Oliva *et al*. 2019; Ahmed *et al*. 2021). Being a water loving plant, rice is highly sensitive to drought stress which significantly affect its grain yield (Shuxing 2014). Drought is becoming a serious yield constraint for various major crops due to global water scarcity (Wattoo *et al*. 2018; Hussain *et al*. 2019; Shokat *et al*. 2020a). A recent study using the yield and metrological data from 1980 to 2015, reported the yield decline up to 21% in wheat (*Triticum aestivum* L.) and 40% in maize (*Zea mays* L.) due to drought on a global scale (Daryanto *et al*. 2016). In rice, mild-drought stress reduced grain yield by 31 to 64%, while severe stress reduced 65–85% yield as compared to normal conditions (Kumar *et al*. 2008). It affects the yield by altering different agronomic and physiological traits including plant height, number of tillers, leaf area, leaf rolling, transpiration rate, accumulation of osmo-protectants, root system and stomatal closure (Nakashima *et al*. 2007; Islam *et al*. 2009; Tong *et al*. 2009). Anthesis stage drought stress can interrupt flowering, floret initiation, (Bajji *et al*. 2002), pollen fertility (Zhou *et al*. 2011) and grain filling, resulting in poor paddy yield. Rice growth is affected by drought at different stages including booting (Shao *et al*. 2014), flowering (Liu *et al*. 2006) and grain filling stage (Zhang *et al*. 2018). However, drought stress at anthesis stage restricts the availability of photosynthates by disturbing the sink capacity (Do *et al*. 2010) and reduces the grain yield, plant biomass and ultimately the harvest index (Blum 2018). It also impairs anther dehiscence, pollen viability and pollen germination in rice resulting in spikelet sterility and more sterile grains in the panicles (Prasad *et al*. 2017). Drought induced spikelet sterility is considered as one of the major causes of yield reduction.

To address the challenge, natural variation in rice germplasm for drought tolerance could be exploited to identify the drought tolerant genotypes, the associated traits and underlying genes (Panda *et al*. 2021). In addition, induced variation via hybridization and mutagenesis could serve as important genetic resource for target breeding (Zafar *et al*. 2020c). For the purpose, scientists have started to put efforts to breed green super rice (GSR), an elite rice type that could withstand multiple stresses with high nutrient use efficiency (Wing *et al*. 2018; Jewel *et al*. 2019). The idea was given by a famous rice geneticist Qifa Zhang in 2007 (Zhang 2007), which was later implemented by a team of International scientists from China and International Rice Research Institute (IRRI), Philippines (Yu *et al*. 2020). The present study was conducted to evaluate 22 selected GSR lines along with 4 local rice cultivars for drought tolerance in Pakistan, and identify agronomic and physiological traits associated with drought tolerance in GSR. In addition, the contrasting drought tolerant and sensitive lines were assessed for gene expression profile to identify underlying genes related to drought tolerance in GSR. This study identified high-yielding drought tolerant GSR lines and provided us knowledge about drought tolerance related traits, and novel drought related genes.

## MATERIALS AND METHODS

### Experimental Site

The field experiment was conducted at National Institute for Genomics and Advanced Biotechnology, NARC, Pakistan (33.684°N and 73.048°E) during rice growing period (May to October, 2020). To minimize the water infiltration from control to drought plot, 6-8 feet path was made between both plots and further, plastic film was applied under the soil surface with depth of 60 cm.

### Experimental design

The 22 diverse GSR lines were selected based on diverse phenotypic characteristics from the 552 GSR genotypes (Table S1). Twenty-two GSR lines along with 4 checks were evaluated using split plot randomized complete block design with two treatments (well-watered and drought) each having three replications. Seeds were sown in nursery trays and thirty days old seedlings were transplanted in the field. Each plot consisted of five rows of 10 plants with 30cm row/row and plant/plant distance (Yugandhar *et al*. 2017). Both plots were irrigated normally (8-10cm) till anthesis stage. Fertilizer, weedicide and insecticide application was done according to recommended dosage. Crop cultivation is carried-out according to normal cultural practices.

Drought was imposed for 30 days by with-holding the applied water at the beginning of anthesis stage. Physiological traits were recorded during last week of stress. After thirty days, field was re-watered. At physiological maturity, five representative plants were selected for the measurement of agronomic traits, from the three middle rows of each replication to avoid confounding border effects (Chaturvedi *et al*. 2017).

### Physiological measurements

#### Cell membrane stability (CMS)

Leaf samples were collected at last week of drought stress to examine the cell membrane stability by recording the electrolyte conductivity using electrical conductivity meter (HI 9811-5 Portable EC meter HANNA® instruments-USA). Flag leaves from three plants per replicate (of each genotype) were collected from both control and drought stress fields in 20ml glass vials. Further measurement was recorded as proposed by (Tripathy *et al*. 2000). CMS was formulated as the reciprocal of cell membrane injury by using following formula (Blum and Ebercon 1981).

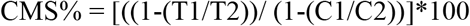

Where, T and C refer to stressed and controlled, respectively. C1(initial control), T1(initial stress) and after autoclave, C2 (final control), T2 (final stress) were the assumed conductance.

#### Normalized difference vegetation index (NDVI)

NDVI is a spectral reflectance-based measure of the density of green vegetation on a land area. NDVI measurements were taken using GreenSeeker™ Handheld Optical Sensor Unit (NTech Industries, Inc., USA), keeping the sensor at 0.5-1 m above the central rows of all the genotypes individually in three replications of both control and stress field plots (Govaerts and Verhulst 2010).

### Pollen fertility test

About five to eight mature spikelets from five panicles (one from each plant) were collected in the morning before anthesis. Spikelets were fixed in FAA solution (formaldehyde: ethanol: acetic acid with a ratio of 1:18:1, respectively) until staining. Anthers were crushed with a forceps on glass slide to release pollens which were immersed in 1% Potassium Iodide (I2-KI) solution followed by observation under the light microscope (NIKON DIGITAL SIGHT DS-Fi2). Pollens which stained black and circular were considered fertile, while those stained red-orange and irregular shape were considered sterile (Zafar *et al*. 2020b). Pollen fertility percentage (PFP) was calculated using the following formula:

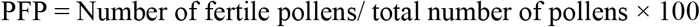

### Measurement of agronomic traits

Agronomic traits including plant height per plant (PH), tillers per plant (TPP), grain yield per plant (GY), straw yield per plant (SY), total biomass per plant (TBM), 1000-grain weight (TGW) and grain length (GL) were recorded manually. Harvest index (HI) was calculated as the ratio of GY to TBM). Drought susceptibility index (DSI) was calculated as ((1-Y/ YP) / D)) as described earlier (Khanna-Chopra and Viswanathan 1999; Zafar *et al*. 2020a). Measurements for these traits was carried-out on five randomly selected plants of each genotype from each replication by following the method (IRRI 2002).

### RNA extraction and cDNA synthesis

Total RNA was extracted from the panicles of selected drought tolerant and sensitive genotypes from both well-watered (WW) and drought-stressed plants. Panicles were harvested from plants and immediately kept in liquid nitrogen followed by storage at -80 C to avoid the denaturation of RNA. The PureLink RNA Mini kit (Thermo Fisher Scientific) was used to extract the total RNA, in accordance with the manufacturer’s protocol. The quality of isolated RNA was observed on 1.5% RNase-free agarose gel, and quantified using the BioSpec-nano spectrophotometer. 1 µg of total RNA was used to reverse transcribe into cDNA using RevertAid Reverse Transcriptase kit, (Thermo-Fisher Scientific) following the manufacturer’s instructions.

### DEGs selection and qRT-PCR

Ten differentially expressed genes (DEGs) under drought stress were selected from a comparative transcriptome study in rice (Huang *et al*. 2014). To our knowledge, these genes have not been studied before specifically for drought response. In addition, we studied expression of three known drought tolerance related genes; *Oryza sativa Salt-, ABA- and Drought-Induced RING Finger Protein 1* (*OsSADR1*) (Park *et al*. 2018), *Drought-Hypersensitive Mutant1* (*DSM1*) (Ning *et al*. 2009), *Drought tolerance 11* (*OsDT11*) (Li *et al*. 2017). Selected genes are listed in Table S2. Coding sequences (CDS) of the selected DEGs were retrieved from the Rice Genome Annotation project (http://rice.plantbiology.msu.edu/cgi-bin/gbrowse/rice/). A gene-specific pair of primers was designed using AmplifX version 1.7.0 software, and primer sequences are listed in Table S3.

Quantitative real time-PCR (qRT-PCR) was carried out to determine the relative expression levels of 13 selected genes on StepOne™ Real-Time PCR System (Thermo Fisher Scientific) using Maxima SYBR Green. The delta cT method was used to calculate the relative expression level of each gene, and rice *Actin1* gene was used to normalize the expression (Fang *et al*. 2021; Zafar *et al*. 2021).

### Statistical analysis

Morpho-physiological traits data were analyzed by analysis of variance using SPSS software according to split plot randomized complete block design. Principal component analysis (PCA) was done through XL-STAT software (ver. 2018) to categorize various physiological and morphological traits (Mohammadi and Prasanna 2003). Pearson’s correlation matrix analysis was done using ‘‘cor’’ package in R studio. The p-values for the coefficient of correlation (r) were obtained by applying Student’s *t-test* with the “cor.test” function in R-studio. In the correlation matrix plot, only significant relationships were labelled with stars. Expression pattern significance were calculated using *t-test*.

## RESULTS

### ANOVA showed significant variation among GSR accessions under drought stress

Analysis of variance (ANOVA) was performed to see the significant differences of variation among the genotypes and water treatments for physiological and yield-related traits. ANOVA showed significant variation (*P*<0.01) among the tested genotypes for PH, GY, HI, TGW, GL and NDVI (Table 1). While non-significant differences were observed for TPP, SY and PFP. There was no significant effect on the studied traits among the replications which strengthen the reliability of this experiment. Drought significantly affected PH, GY, SY, HI, GL, PFP and NDVI while traits like TPP, TBM and TGW were not affected by drought. The genotype × environment interaction was also significant for PH, TPP, SY, HI, TGW, PFP and NDVI (Table 1) where pronounced reduction was recorded under drought conditions. Since PH, GY, HI, GL, and NDVI displayed significant difference for genotypes as well as drought treatment, these traits could be key selection markers for drought tolerance screening in rice.

**Table 1.** Mean square values from the analysis of variance for the effect of genotype, environment and their interaction on agronomic and physiological traits.

| Source of variation | DF | PH | TPP | GY | SY | TBM | HI | TGW | GL | PFP | NDVI |
| --- | --- | --- | --- | --- | --- | --- | --- | --- | --- | --- | --- |
| **Genotype** | 25 | 687.55** | 47.97<sup>ns</sup> | 956.25** | 599.24<sup>ns</sup> | 2136.97<sup>ns</sup> | 0.0081* | 42.40** | 4.02** | 727.03<sup>ns</sup> | 0.0042** |
| **Replication** | 2 | 42.40<sup>ns</sup> | 107.97<sup>ns</sup> | 700.96<sup>ns</sup> | 875.40<sup>ns</sup> | 3633.82<sup>ns</sup> | 0.00056<sup>ns</sup> | 8.87<sup>ns</sup> | 0.38<sup>ns</sup> | 15.07<sup>ns</sup> | 0.02<sup>ns</sup> |
| **Treatment** | 1 | 1243.68** | 0.27<sup>ns</sup> | 19649.7* | 5543.01* | 2969.05<sup>ns</sup> | 0.36** | 106.91<sup>ns</sup> | 6.31** | 5434.18* | 0.14** |
| **Genotype × Treatment** | 25 | 57.59** | 37.34** | 236.93<sup>ns</sup> | 520.24* | 1029.61<sup>ns</sup> | 0.0033** | 3.72** | 0.20<sup>ns</sup> | 771.38** | 0.0013* |

Significance levels are indicated: \*, $P < 0.05$ ; \*\*, $P < 0.01$ . df, degrees of freedom; PH, plant height; TTP, tillers per plant; GY, grain yield; SY, straw yield; TBM, total biomass; HI, harvest index; TGW, thousand-grain weight; GL, grain length; PFP, panicle fertility percentage; NDVI, normalized difference vegetation index.

### Principal component analysis revealed genetic variation among GSR accessions under drought

Separated PCA analyses were performed to develop a trait–genotype (T– G) biplot and to detect genetic variation among the studied genotypes for various morpho-physiological traits under well-watered and drought-stressed conditions. Under WW environment, a biplot was drawn between PCI and PC2 explaining31.3% and 25.6%, of total variation, respectively (Figure 1a). Our results indicate SY, GY, HI and TBM were in the opposite direction to NDVI, TGW and GL indicating their opposite relationship with each other. In addition, the GSR lines were mostly clustered near the origin and show less genetic variability, while checks Kashmir Basmati, Kissan Basmati, and IR-64 were widely distributed apart from the origin and showed remarkable genetic variability (Figure 1a). In case of drought treatment, the PC1 alone accounted for 51.10% of the total variability, while PC2 shared 15.20% (Figure 1b). Results of this experiment shows GY were clustered closer to PFP, CMS, TBM and HI while it was in opposite direction to NDVI and DSI. In contrast to the WW treatment, many GSR lines namely NGSR-3, NGSR-15, NGSR-18, NGSR-13, NGSR-21 and NIAB-IR-9 fall near the apex of the biplot and show remarkable genetic variation under drought stress (Figure 1b). The check varieties Kissan Basmati, NIAB-IR-9, and Kashmir Basmati also showed considerable genetic variability and reputation of these accessions for further selection in breeding programs.

**Figure 1.**
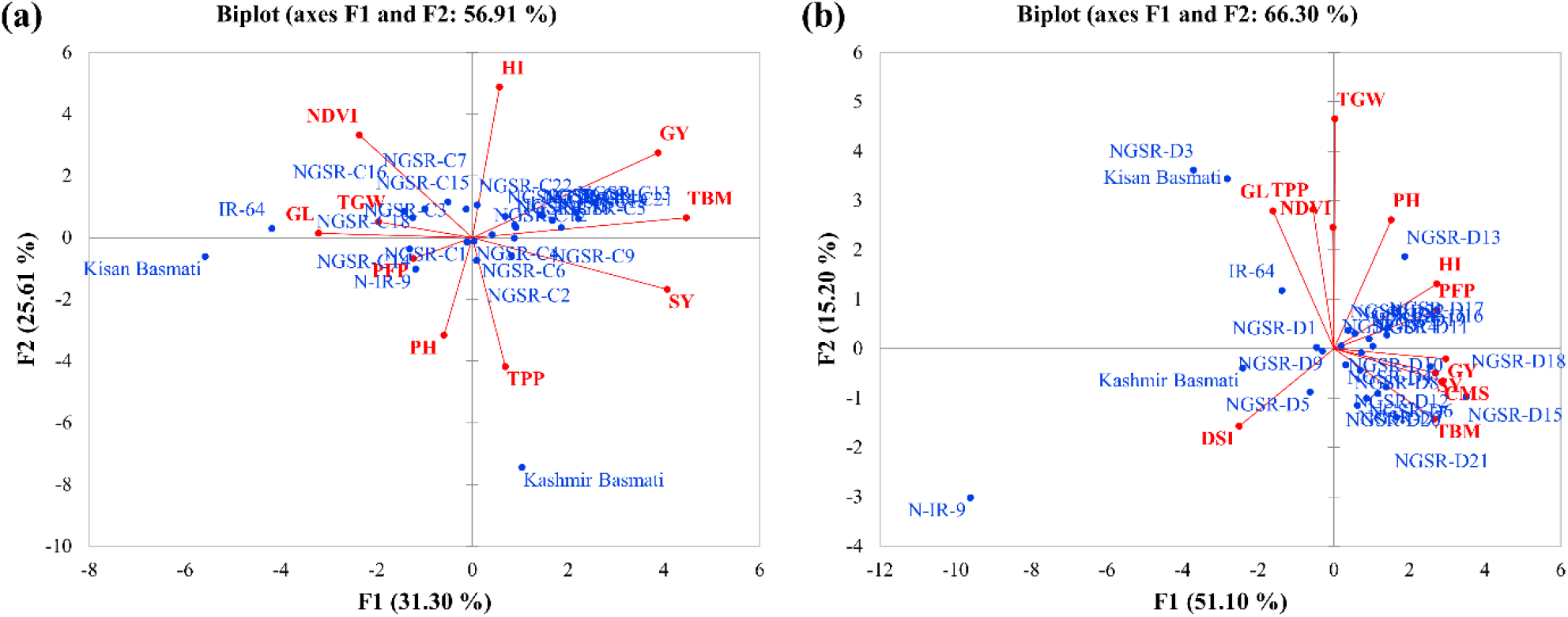
PCA showing biplot for genotypes and studied traits (a) under normal condition and (b) under drought stress condition

**Figure 1.**
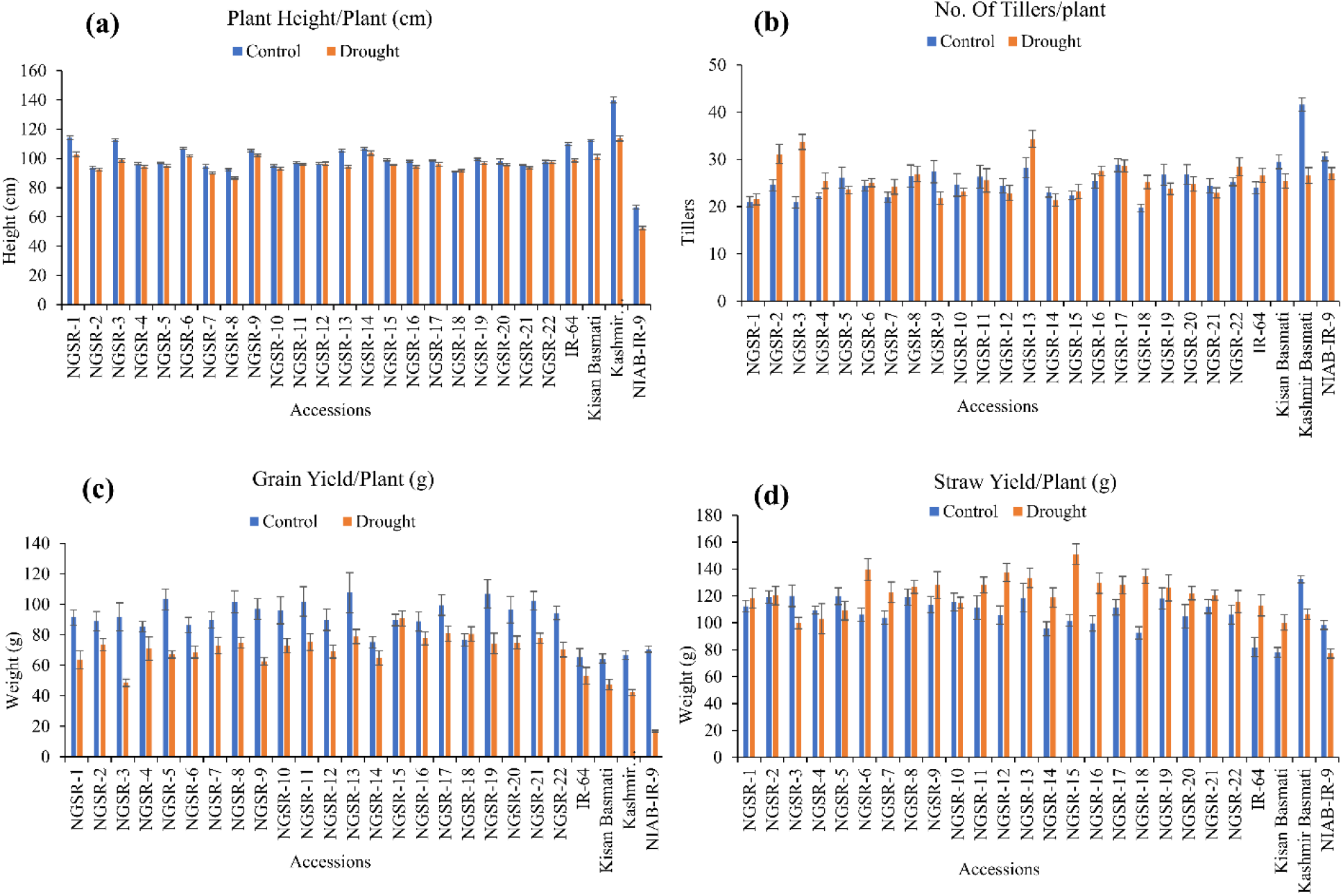
Effect of drought stress on (a) plant height/plant, (b) tillers/plant, (c) grain yield/plant and (d) straw yield/plant. Values are means ± SD.

### Mean performance of GSR accessions for studied traits

Drought stress showed a remarkable reduction in grain yield and yield-related traits in all studied genotypes except NGSR-15 and NGSR-18 (Figures 2-3). Among the 22 GSR lines, the minimum PH (86.6cm) under drought condition was attained by NGSR-8 and the maximum (103.7cm) was recorded by NGSR-14. Whereas, among 4 checks the maximum PH was recorded for Kashmir Basmati (113.6cm) and the minimum was depicted by NIAB-IR-9 (52.3cm). All GSR lines demonstrated higher PH than drought sensitive check NIAB-IR-9 (Fig. 2a). These results suggested that the GSR lines were comparatively less affected by drought stress and maintained the normal plant growth.

**Figure 2.**
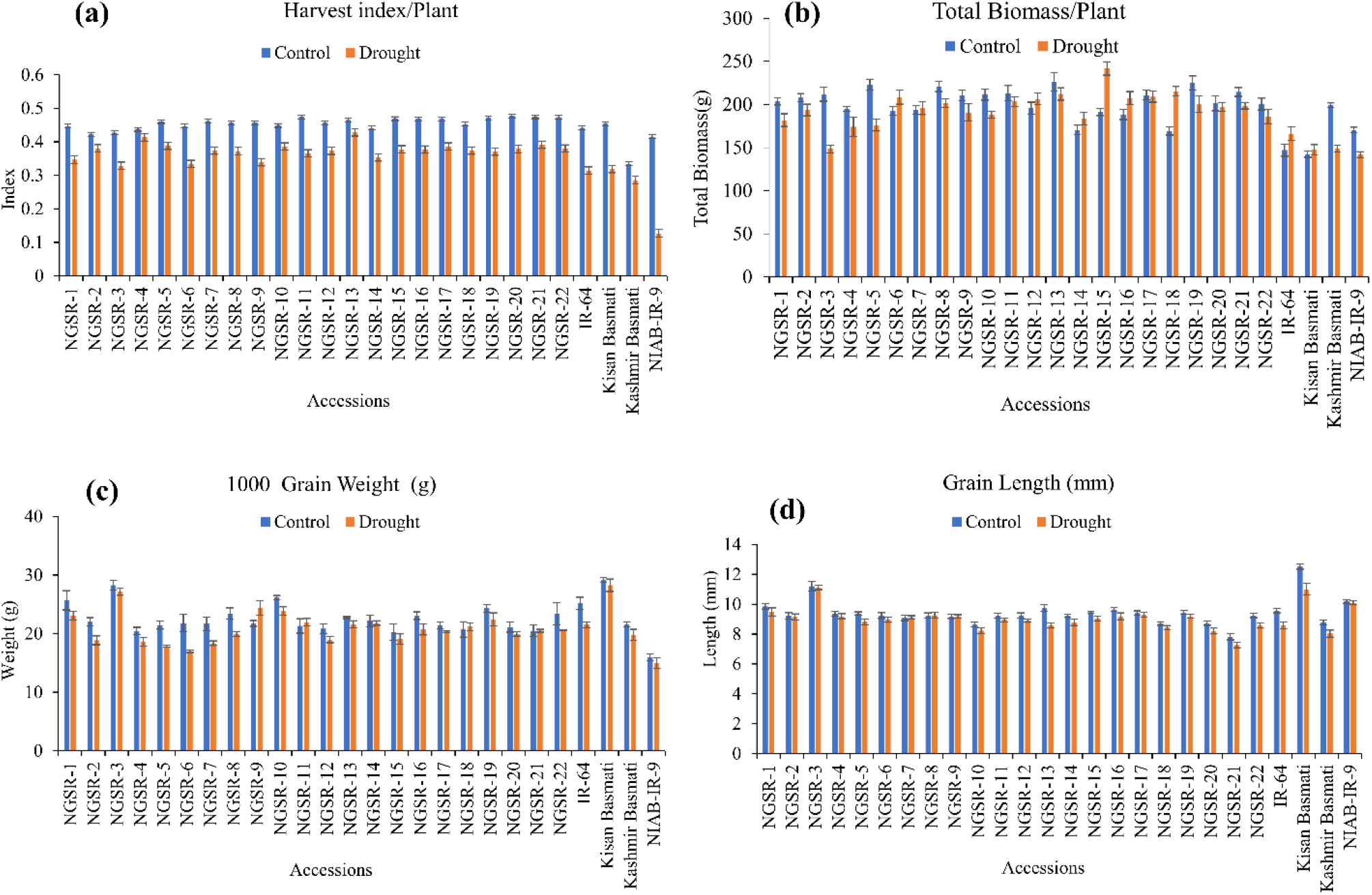
Effect of drought stress on (a) harvest index/plant, (b) total biomass/plant, (c) 1000-grain weight and (d) grain length. Values are means ± SD.

Overall, TPP were not significantly affected under drought stress except for Kashmir basmati, while few of the GSR lines showed increased TPP under drought (Figure 2b).GY is the most important agronomic trait of economic importance, and drought stress affected the GY in most genotypes except NGSR-15 and NGSR-18 (Figure 2c). Under drought stress, maximum GY was reported by NGSR-15 (90.7g), and the minimum by the NGSR-3 (48.6g). Among the checks, IR-64 showed the maximum GY^-1^ (53.1g) whereas the minimum showed by the drought sensitive NIAB-IR-9 (16.8g). All the GSR lines (except NGSR-3) demonstrated higher grain yield than the check varieties, even the drought sensitive NGSR-3 accounted for higher grain yield than the sensitive check NIAB-IR-9 (Fig. 2c). These results suggested that the grain yield of GSR lines was less affected by drought stress as compared to local checks.

SY was generally increased in most GSR accessions along with two check varieties IR-64 and kisan basmati under drought stress (Figure 2d). The maximum increase in SY was observed in NGSR-15, NGSR-6, NGSR18 and NGSR-12. However, NGSR-3, Kashmir basmati and NIAB-IR-9 showed a decrease in SY under drought. It is noteworthy that SY was only decreased in the most drought sensitive GSR line and checks, thus it is considered an important trait for drought escape at flowering stage. This is because plants tend to continue their vegetative stage bypassing the flowering stage until they got favorable conditions.

Generally, drought stress negatively impacted the HI in all genotypes, but the non-GSR lines showed higher decrease compared to GSR lines with the highest decrease observed in our drought sensitive check NIAB-IR-9 (Figure 3a). These results suggest that GSR lines have the potential to maintain the HI under drought stress conditions (Fig. 2a).

**Figure 3.**
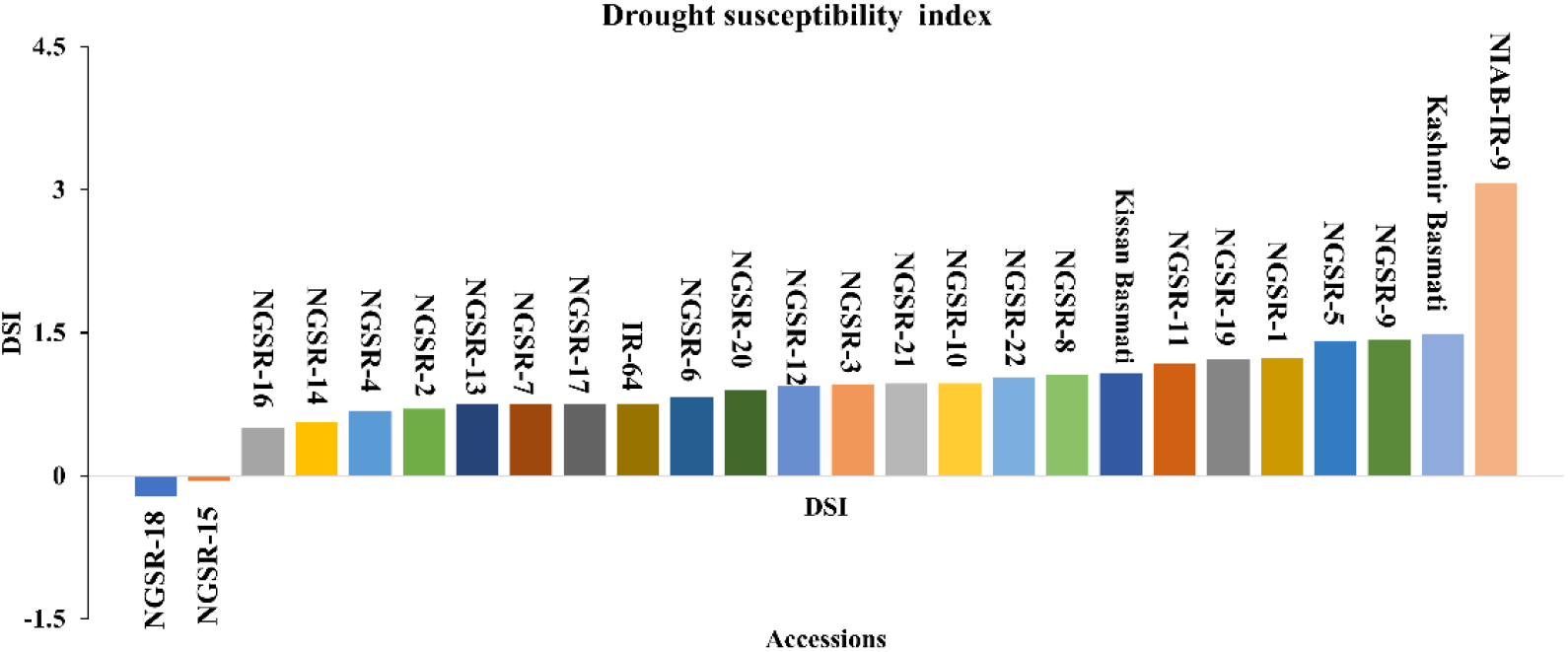
Frequency distribution of drought susceptibility index for grain yield showing the degree of susceptibility to drought stress. Genotypes below the line are declared as most drought tolerant genotypes.

TBM was not significantly affected under drought stress except for Kashmir basmati and NIAB-IR-9 (Figure 3b). Similarly, TGW was not significantly affected under drought stress in tested genotypes (Figure 3c).

Drought stress had a significant effect on GL, however, differences for genotypes were non-significant (Figure 3d). Three genotypes including NGSR-3, NGSR-1 and NGSR-15, showed longer GL. The genotypes NGSR-3 and NGSR-21 showed the highest (11.1 mm) and the lowest (7.3 mm) GL, respectively. Among the experimental checks, the maximum (10.9 mm) and the minimum (8.03 mm) GL depicted by Kissan Basmati and Kashmir basmati, respectively. Overall, GSR lines maintained the grain length under drought stress as compared to sensitive checks (Fig. 3d), except NGSR-21 which showed the reduced grain length (7.3mm).

### Drought susceptibility index

DSI is an important indicator of drought tolerance and lower value indicates better tolerance. Overall, the GSR lines showed lower DSI compared to local varieties, and the genotypes NGSR-18, NGSR-15, and NGSR-16 exhibited the lowest DSI among 22 GSR lines, showing their potential towards drought tolerance (Figure 4). In contrast, highest DSI was recorded for NIAB-IR-9 followed by Kashmir basmati, indicating the least drought tolerance among tested genotypes. Overall, these results demonstrated that GSR lines generally performed better than the checks.

**Figure 4.**
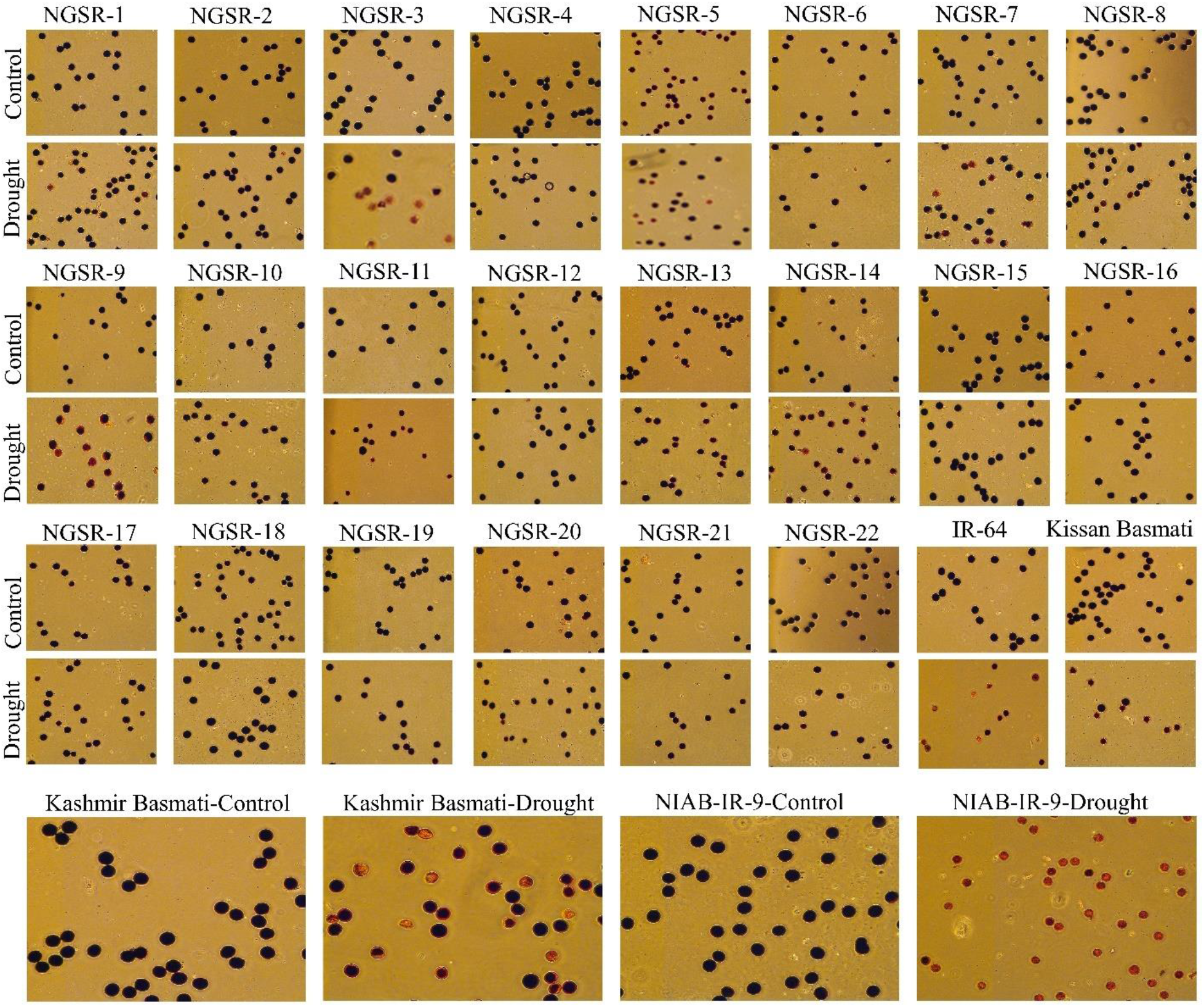
Examination of pollen fertility of the 22 GSR lines and 4 checks with I_2_-KI solution staining of the mature pollen grains. The sterile pollen grains failed to be stained or stained weakly, indicating that they did not contain starch or contained irregularly distributed starch, whereas the viable pollen grains were stained deep brown.

### Pollen fertility percentage

Pollen fertility is an important indicator of drought tolerance as it directly affects the seed setting and ultimately the grain yield. The microscopic analyses of potassium-iodide (I_2-_KI) stained anthers revealed significant differences in PFP between tolerant and sensitive genotypes. Overall, the GSR lines maintained higher PFP under drought stress compared to non-GSR checks (Figure 5). While most of the GSR lines showed completely fertile pollens under drought a higher sterility up to 57.1% was recorded in NGSR-3 (Figure 5, 6a). In contrast, check varieties except the kisan basmati showed lower PFP as compared to GSR under drought whereNIAB-IR-9 showing only 3.6% PFP (Figure 6a). These findings suggest that PFP could be a good indicator for drought tolerance in rice.

**Figure 5.**
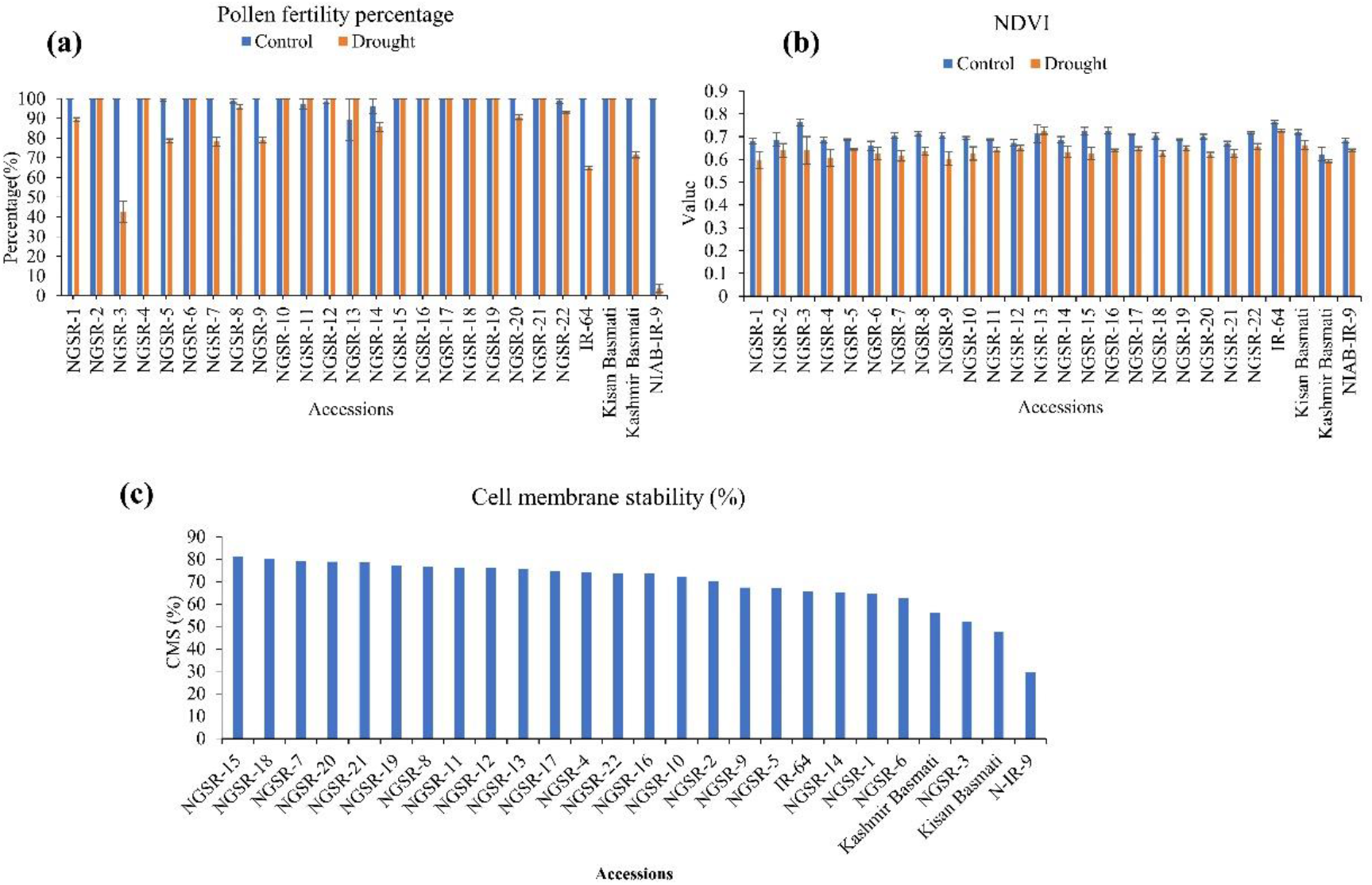
Effect of drought stress on (a) pollen fertility percentage, (b) NDVI, and (c) cell membrane stability-CMS. Values (except CMS) are means ± SD.

**Figure 6.**
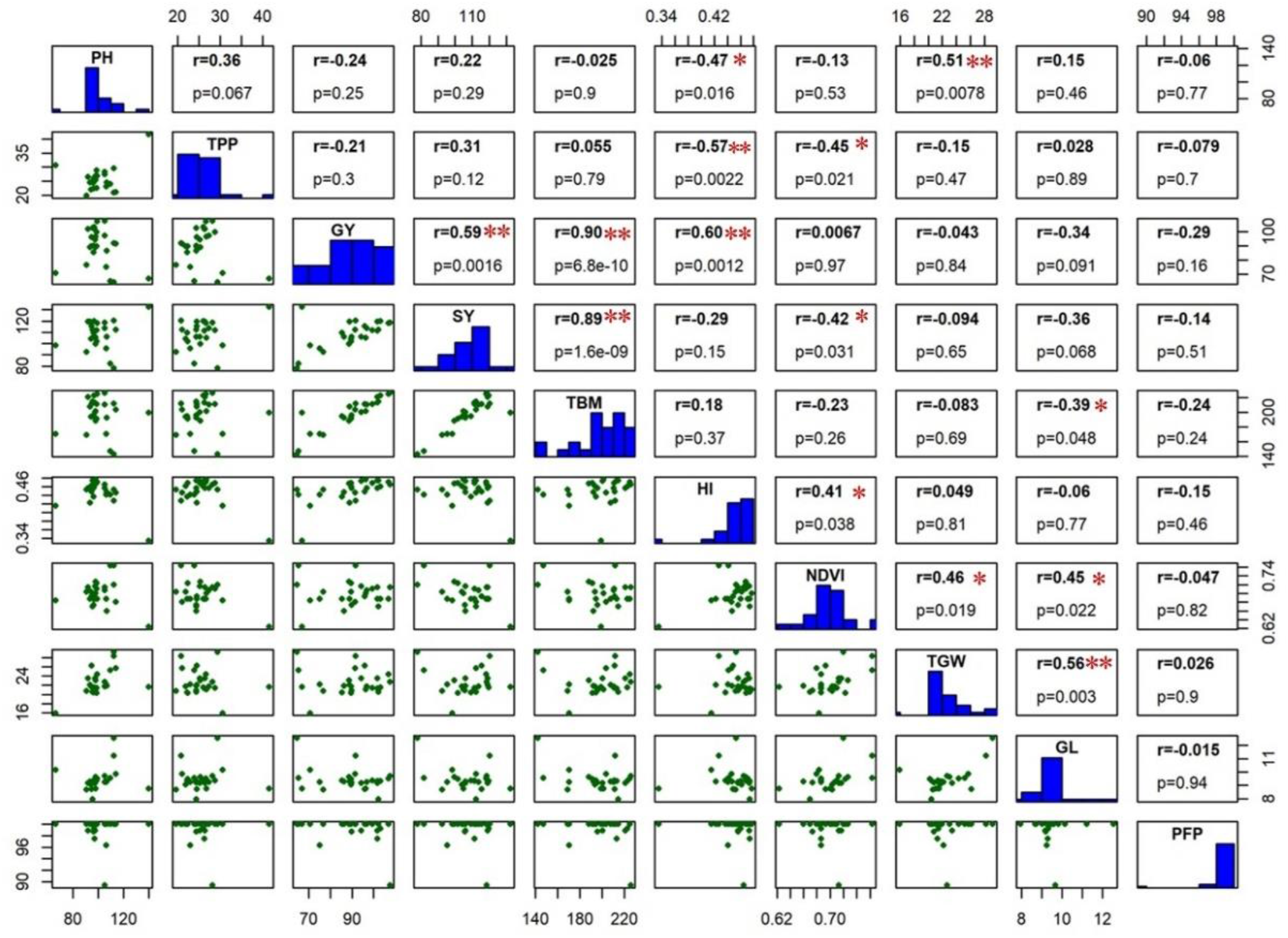
The scatter-matrix below the histogram, and correlation co-efficient value with p-value above the histogram calculated from means of all the studied traits under WW environmental condition. The p-value of all correlations was 0.05* and 0.01**.

### NDVI

NDVI is recently emerged as an indicator of plant health. We observed a considerable decrease in the value of NDVI under drought stress for GSR and check varieties (Figure 6b). The genotypes NGSR-13, IR-64 and Kissan Basmati showed the highest NDVI (>0.7) under drought. Among GSR lines the maximum NDVI (0.7) was reported in NGSR-13, whereas the minimum (0.59) in NGSR-1. Similarly, among the check verities, the highest NDVI depicted by IR-64 (0.72) whereas, minimum by Kashmir Basmati (0.6) (Fig. 6b). Since drought often causes leaf yellowing in plants, the reduced NDVI values under drought could be associated with yellow leaves.

### Cell membrane stability

CMS indicates the stress tolerance ability of plant cells. Again, GSR lines showed higher CMS% than non-GSR where NGSR-15 showed highest (81.1%) CMS followed by NGSR-18 (Figure 6c) while lowest was measured in NIAB-IR-9 (29.5%).

### Correlation of grain yield with other agronomic traits

Understanding the correlation of grain yield with other agronomic and physiological traits is of prime importance as it helps to identify certain pre-breeding traits that could be best indicators of grain yield. Under WW environment, GY has shown a significant positive correlation with SY (r = 0.59**), TBM (r = 0.90**) and HI (r = 0.60**) (Figure 7).

**Figure 7.**
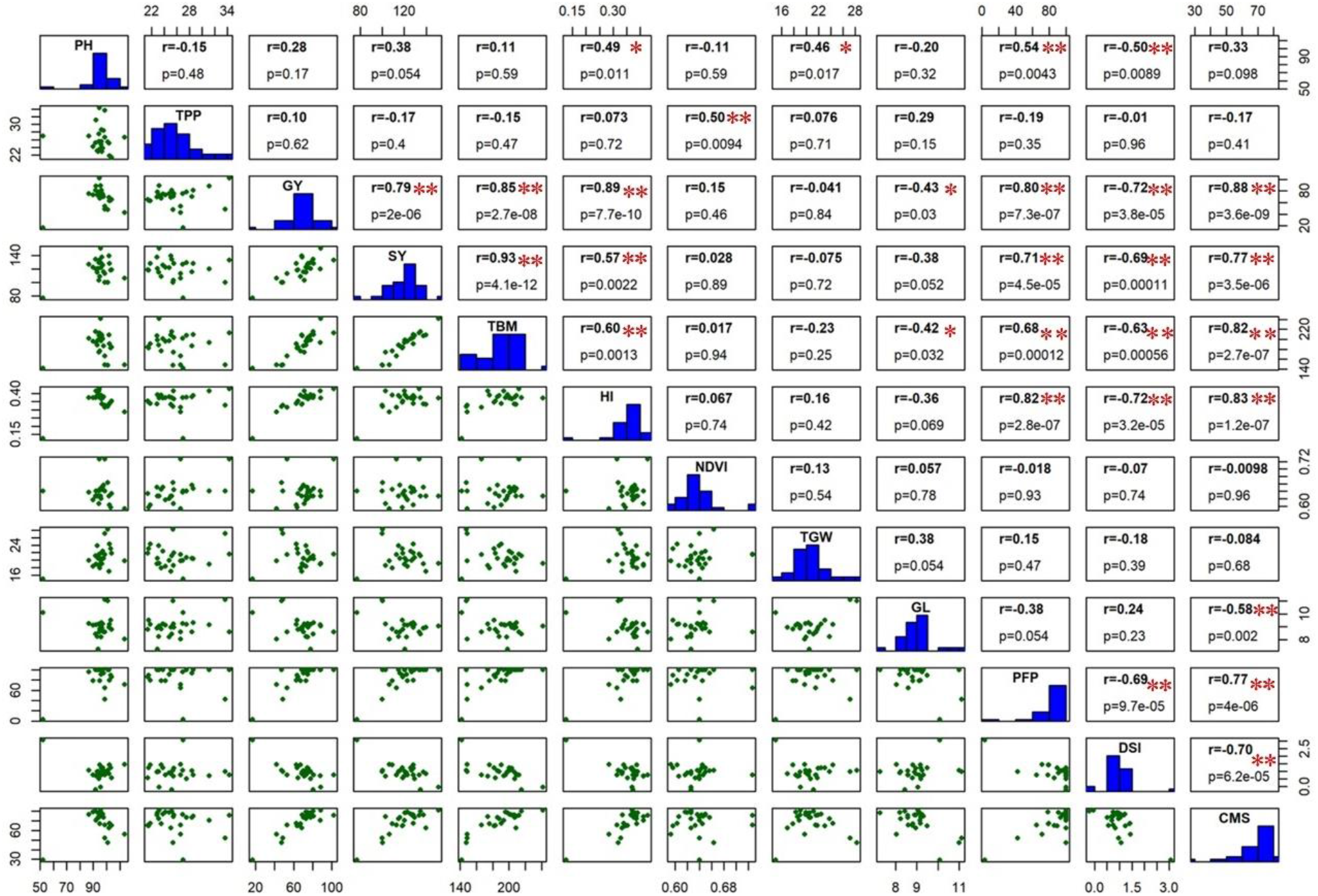
The scatter-matrix below the histogram, and correlation co-efficient value with p-value above the histogram calculated from means of all the studied traits under drought stress condition. The p-value of all correlations was 0.05* and 0.01**.

In addition, a significant negative correlation was found for TPP with HI (r = -0.57**), which suggest the importance of optimum number of tillers for better HI and GY.

Under drought stress, GY has shown a significant positive correlation with HI (r = 0.89***), CMS (r = 0.88***), TBM (r = 0.85***), PFP (r = 0.80***), and SY (r = 0.79***), while a significant negative correlation was found with DSI (r = -0.72***) and GL (r = -0.43*) (Figure 8). In addition to the GY, PFP has shown a significant positive correlation with HI (r = 0.82**), CMS (r = 0.77**), SY (r = 0.71**), and TBM (r = 0.68**), while a significant negative correlation of PFP was observed with DSI (r = -0.69**). Notably, DSI had significant negative correlations with GY (r = -0.72**), HI (r = -0.72**), CMS (r = -0.70**), SY (r = -0.69**), PFP (r = -0.69**), TBM (r = -0.63**), and PH (r = -0.50**), which suggest the importance of DSI being an important indicator of drought susceptibility in rice (Figure 7). These findings revealed important agronomic and physiological traits to be considered as reliable selection criteria for screening rice germplasm against drought tolerance.

### Expression analysis of drought related genes

To see the role of drought responsive genes in drought tolerance of GSR, we analyzed the expression pattern in a selected drought tolerant and drought sensitive genotypes using quantitative real-time PCR (Figure 9). The genotype NGSR-15 was selected as drought tolerant, and NGSR-3 and NIAB-IR-9 were chosen as drought susceptible genotypes (Figure 10). Ten differentially expressed genes (DEGs) were selected for qRT-PCR analysis from the comparative transcriptome dataset between drought sensitive (HHZ) and tolerant (H471) genotypes (Huang et al. 2014). These genes have not been studied previously for their role in drought tolerance, except the transcriptome analysis. In addition, we analysed expression of 3 previously characterized genes for drought tolerance in rice (*OsSADRI, OsDSM1*, and *OsDT11*).

**Figure 9.**
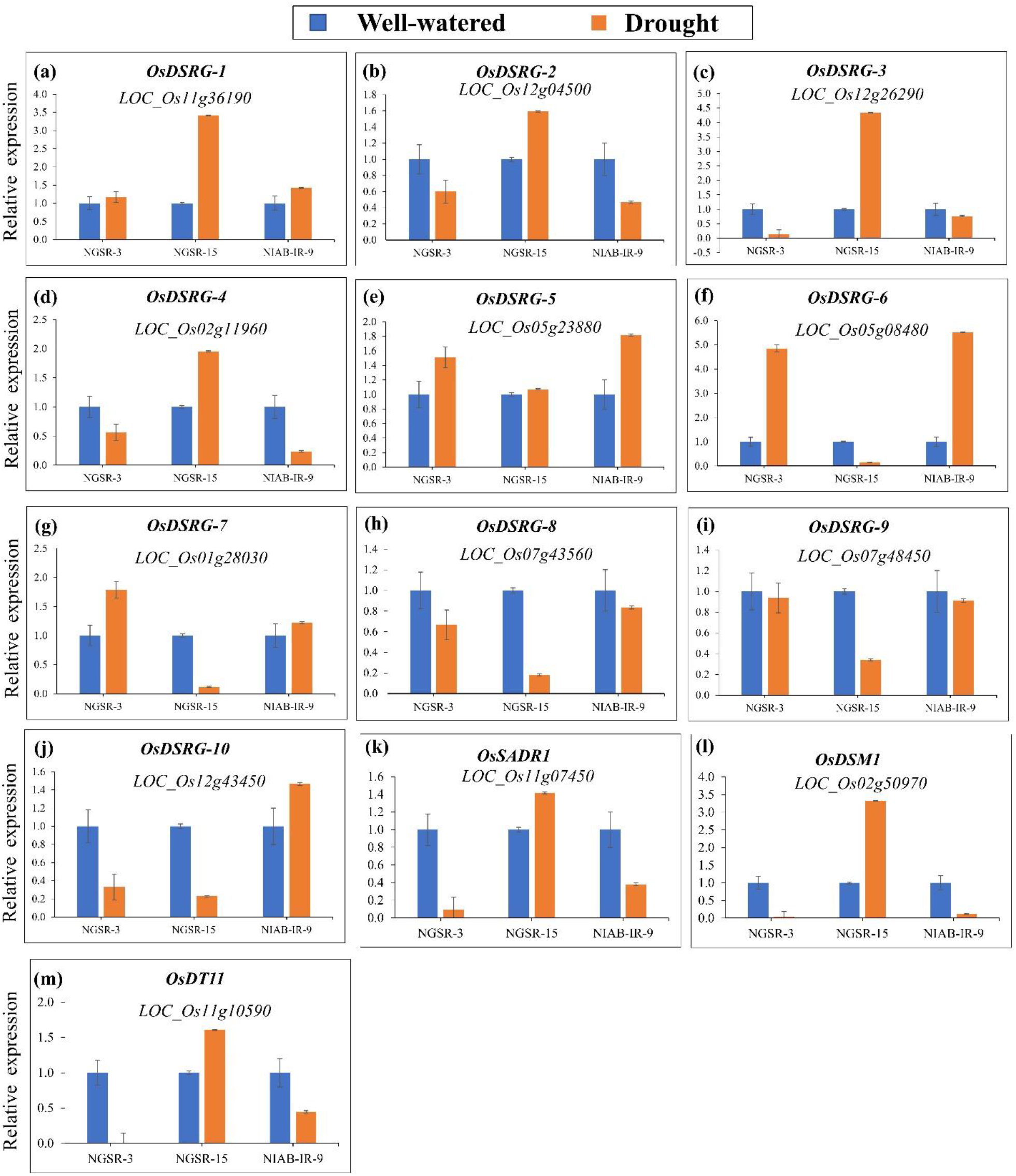
QRT-PCR analysis of *OsDSRG* gene family members in NGSR-3, NGSR-15, and NIAB-IR-9 revealed the relative expression in terms of fold change (log2FC). Young penicle tissues (∼ 1.5 cm) of 3 selected genotypes of were employed in this analysis. Rice actin gene (*OsACT1*) was the internal control gene. Values of 3 technical replicates (n = 3) were expressed as mean ± SD.

**Figure 10.**
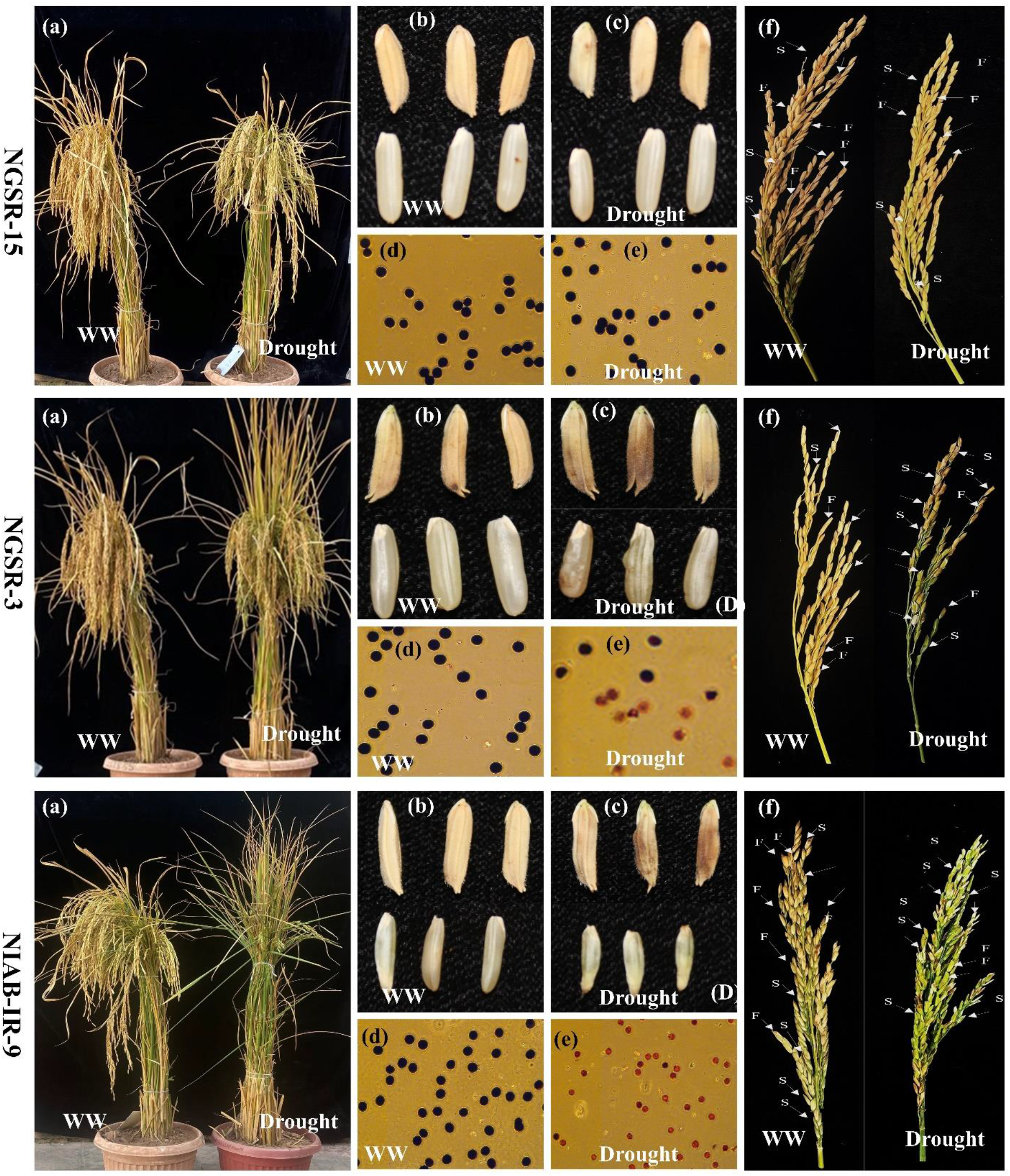
Phenotypic comparison of plants, grain length and shape (husked & de-husked), pollen viability, and panicle fertility of NGSR-15, NGSR-3 and NIAB-IR-9 under well-watered (WW) and drought stress. Abbreviations: F, fertile spikelets; S, sterile spikelets. Spikelets with open tips represent sterile spikelets with no seed set.

Our results indicated that four genes (*LOC_Os11g36190, LOC_Os12g04500, LOC_Os12g26290* and *LOC_Os02g11960)* were upregulated in drought tolerant genotype (NGSR-15) and downregulated in drought sensitive genotypes (NGSR-3 and NIAB-IR-9) (Figure 9). This suggest that these genes may positively regulate drought tolerance in rice. Three genes (*LOC_Os05g23880, LOC_Os05g08480* and *LOC_Os01g28030)* were downregulated in NGSR-15, while upregulated in drought sensitive genotypes (NGSR-3 and NIAB-IR-9), suggesting a negative regulation for drought tolerance (Figure 9). Two genes *LOC_Os07g43560* and *LOC_Os07g48450* were downregulated under drought stress in all genotypes, and thus may not be related to drought tolerance (Figure 9). The gene *LOC_Os12g43450* was upregulated in NIAB-IR-9, while downregulated in both NGSR-3 and NGSR-15, suggesting that this gene might be related to non-GSR rice. These results revealed a differential expression pattern of genes among drought tolerant and sensitive genotypes, and thus could be employed for molecular identification of drought tolerant rice genotypes at large scale. Notably, we observed an increased expression of previously known drought tolerance related genes (*OsSADRI, OsDSM1*, and *OsDT11*) in NGSR-15, while opposite was observed for NGSR-3 and NIAB-IR-9, which clearly indicated the role of these genes in drought tolerance.

## DISCUSSION

Rice being a prime diet of fifty percent global population and staple food of many countries is an important grain crop. However, growing rice requires high delta of water where limited water conditions affect its growth and grain yield. Water stress at anthesis directly affects seed setting and grain filling and, ultimately the grain yield (Shokat *et al*. 2020a; Shokat *et al*. 2020b). Green super rice (GSR) was developed by combining best global germplasm and has the potential to maintain the optimum grain yield under different stress conditions (Jewel *et al*. 2019). Moreover, this germplasm has never been evaluated for pre-anthesis drought stress. In current experiment, 22 GSR genotypes and four local lines of Pakistan were used to understand the mechanism of yield reduction at pre-anthesis stages of drought stress. This germplasm was characterized for different agro-physiological traits and then most diverse genotypes were further evaluated by novel drought responsive genes. Yield related traits are important indication of final grain yield (Zafar *et al*. 2020a; Waqas *et al*. 2021). Studies reported that plant genotypes maintained higher plant biomass under drought stress conditions often maintain higher grain number or weight and ultimately the grain yield (Shokat *et al*. 2021). In the current study, we identified NGSR-15 as drought tolerant line as it maintained higher CMS, PFP, TBM, and GY. In contrast, NGSR-3 and NIAB-IR-9 were ranked drought sensitive lines since they showed significant reductions in GY probably due to reduced CMS, PFP, and TBM. A phenotypic presentation of the performance of these genotypes under drought stress is shown in Figure 10. Higher biomass is usually linked with higher photosynthetic rate of the genotypes (Morinaka *et al*. 2006). Our results indicate that biomass partitioning towards grain filling was limited due to flowering stage drought stress and there could be the possibility of that GSR-15 has maintained higher grain yield due to better seed setting under moisture stress conditions. To explain the possible mechanism of higher and lower grain yield for the genotype GSR-15 and NIAB-IR-9 respectively, we associated yield data with few parameters of physiology to understand the physiological basis of yield reduction at flowering stage drought stress.

DSI indicates the extent of susceptibility by drought stress in terms of economically important traits particularly the grain yield. In this study, genotypes NGSR-15 and NGSR-18 showed the lowest susceptibility with value of -0.04 and -0.2 respectively whereas NIAB-IR-9 (check) showed the highest DSI value of 3.6 (Fig 4). Under drought stress, permeability of membranes and leakage of ions occurs from the weak or unstable membranes (Bajji et al. 2002). Likewise, seed setting is dependent on the viability of pollen while limited water availability at flowering stage can cause pollen abortion in sensitive genotypes (Mehri *et al*. 2020). In contrast, plant genotypes show better cell membrane stability (CMS) or pollen fertility could perform better under flowering stage drought stress. In current experiment, better cell membrane stability and pollen fertile percentage (PFP) was exhibited by genotype NGSR-15 while lowest value was recorded for NGSR-3 and NIAB-IR-9 indicating the physiological basis of drought tolerance and drought susceptibility of these genotypes respectively. A correlation and PCA was drawn to test the significance of these parameters in relation to yield and yield related traits and we find a strong significant and positive correlation of CMS and PFP with grain yield (Figure 1b). In contrast, association of grain yield was significant but negative with DSI (Figure 1b) indicating these traits could be selected as pre-breeding traits for flowering stage drought stress in rice. To understand the molecular basis of drought tolerance these three genotypes were further tested through gene expression.

Stress conditions change the expressions of the stress-induced regulatory or effector genes which usually involve in regulation of normal processes of the plants (Ouyang *et al*. 2010). We investigated different categories of DEGs, controlling drought tolerance and sensitivity by up/down regulation of DEGs. Further, this analysis relied on two GSR genotypes and one locally developed genotype, NIAB-IR-9, in order provide an accurate estimate of expression by comparing GSRs with traditional cultivars under flowering stage drought stress. In the agreement with published literature, our expression results suggest the involvement DEGs in drought tolerance or sensitivity (Chen *et al*. 2009; Narsai *et al*. 2013; Zhang *et al*. 2015b). Additionally, another gene, leucine-rich-RLKs, plays a key role in regulation of plant growth under various abiotic stresses and gene; *LOC_Os11g36190* (a leucine-rich receptor-like kinase) is predicted to up-regulated for bacterial leaf blight in rice (Zhang *et al*. 2015a; Ahsan *et al*. 2019). *LOC_Os12g04500* and *LOC_Os12g26290* are also reported as core of jasmonic acid (JA) signaling pathway and in the current experiment, expression of these two genes was increased significantly under prolonged drought period. Moreover, JA signaling genes are also reported to be involved under critical phases of drought stress (Du *et al*. 2007). We found four genes *i*.*e*., *LOC_Os11g36190, LOC_*Os12g04500, LOC_Os12g26290 and LOC_Os02g119600 were upregulated in drought tolerant genotype (NGSR-15) and downregulated in drought sensitive genotypes (NGSR-3 and NIAB-IR-9) indicating their positive relationship with drought tolerance.

Likewise, an increased expression of previously known drought tolerance related genes (*OsSADRI, OsDSM1*, and *OsDT11*) (Ning *et al*. 2009; Li *et al*. 2017; Park *et al*. 2018) in NGSR-15, while an opposite trend was observed for NGSR-3 and NIAB-IR-9. Apart from gene expression, these genotypes also showed a contrast for PFP, CMS and DSI along with clear differences in grain yield suggesting their role at terminal stage drought tolerance.

Through this study we identified molecular and physiological basis of higher grain yield at flowering stage drought stress and the role of OsDSRG family in this tolerance. Importantly, various morpho-physiological traits (PFP,CMS,DSI and HI) had strong association with genes of *LOC* and OsDSRG family and ultimately the grain yield indicating these parameters could be used as pre-breeding triats for drought tolerance. Our results also indicate genotype NGSR-15 was the most drought tolerant while NGSR-3 and NIAB-IR-9 were the most sensitive genotypes. These genotypes can further be used improve rice yield under drought stress, however, in depth mechanism is required to confirm our findings.

## Authors contribution

SAZ, MKN and MRK designed the study. HA, SAZ and MKN planted the fields and collected data. SAZ, MKN and SS analysed the data. HA, SAZ and MKN wrote the manuscript. SS, SI, GMA and MRK provided technical assistance. ASN, JX and ZL developed the GSR material. MRK supervised the whole study and provided technical and financial support.

## Notes

### Competing Interest Statement

The authors have declared no competing interest.

